## Supplementary Information for "ATAC-seq with unique molecular identifiers improves quantification and footprinting"

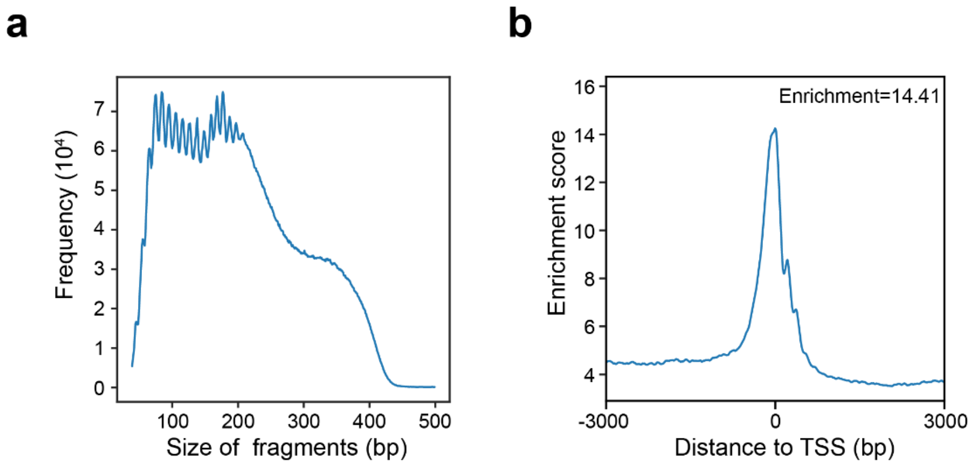


**Supplementary Figure 1: The quality control results of UMI-ATAC-seq**

(a) Distribution of fragment sizes for sample C019 from aligned, filtered and deduplicated reads. (b) Enrichment of ATAC-seq accessibility near Transcription Start Sites (TSS) in sample C019.

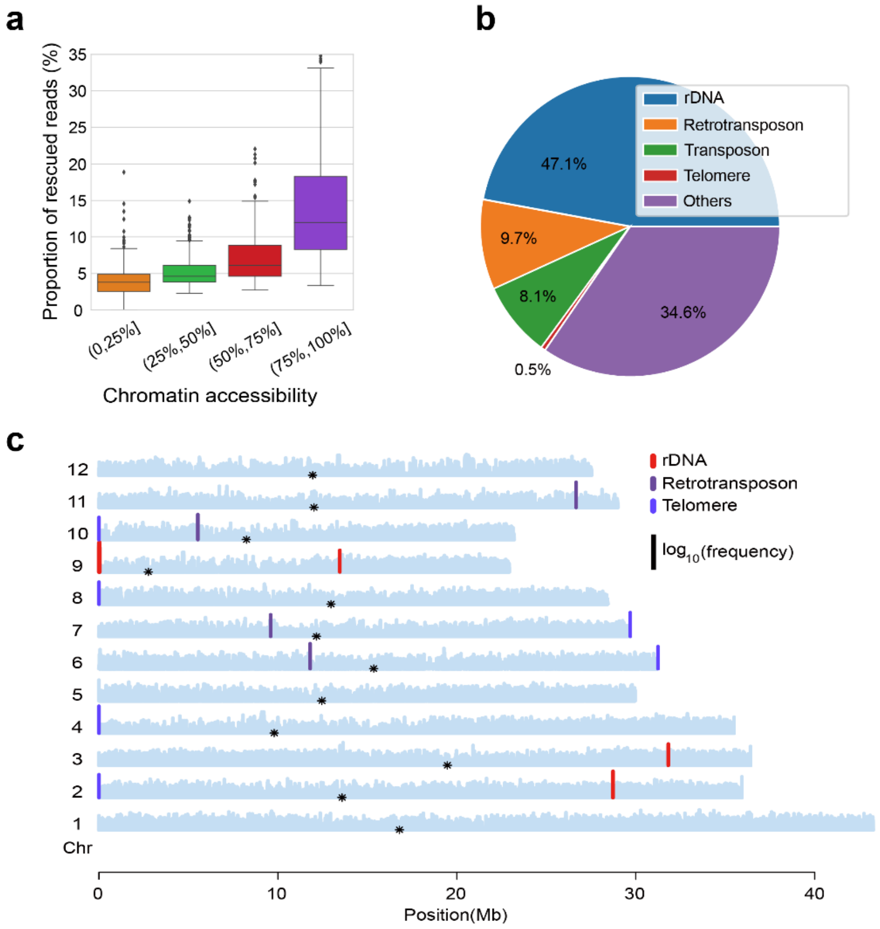


**Supplementary Figure 2: The characteristics of rescued reads which have identical mapping coordinates but different UMI in sample C019.** (a) Box plot illustrating the distribution of rescued reads with different chromatin accessibility. We divided the genome into 1 kb bins and counted the rescued reads in each bin. Then we grouped all bins by 25%, 50% and 75% quantiles. Only reads with mapping quality greater than thirty were used. (b) The annotations of rescued reads. (c) The distribution of rescued reads in the genome. The maximum value of the frequency in a bin (1 kb) is 39,912. Bins annotated as rDNA, retrotransposon and telomere are marked. Black points represent the centromere.

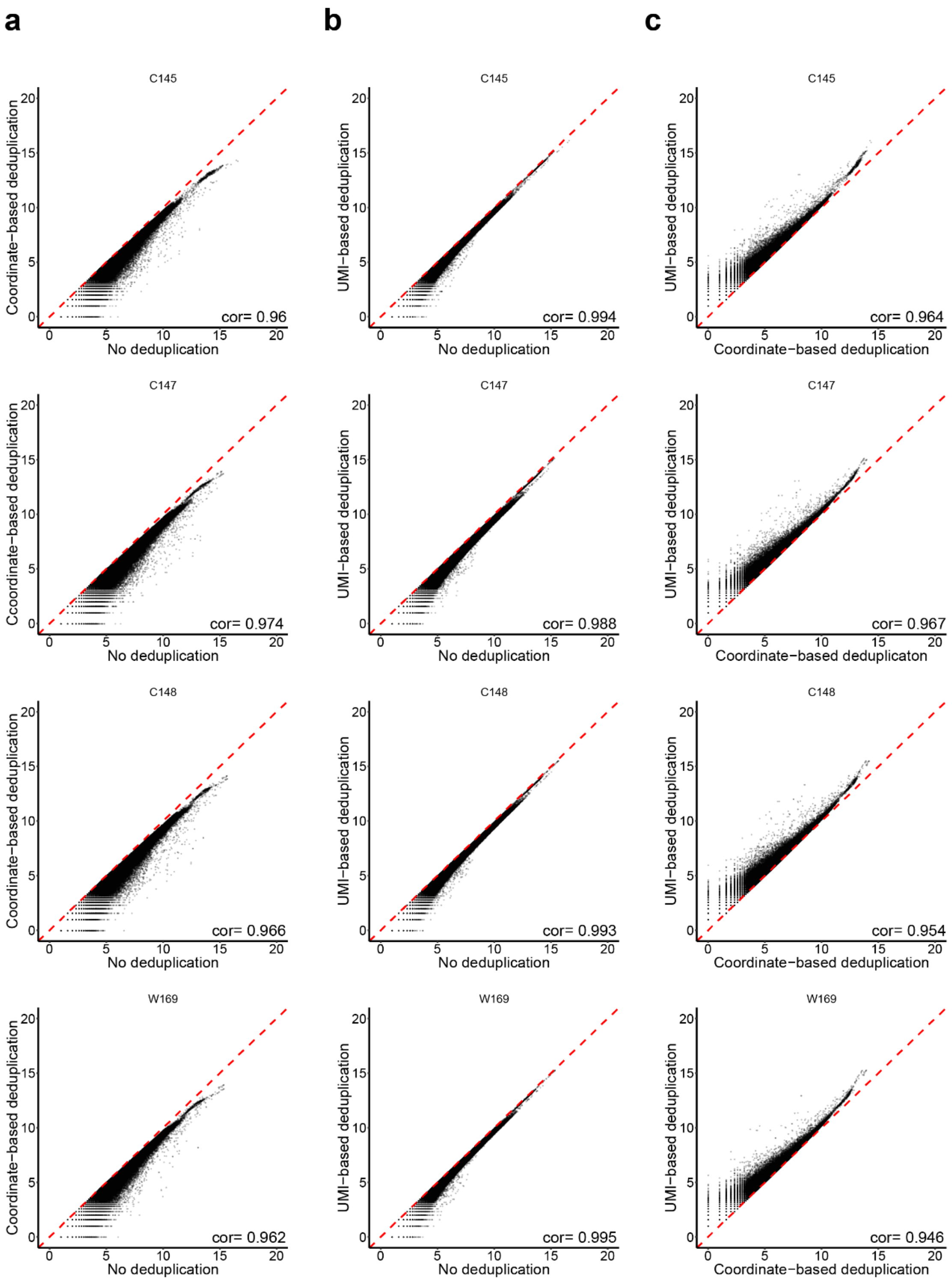
a
b
c
C145
Coordinate−based deduplication
No deduplication
cor= 0.96
C145
UMI−based deduplication
No deduplication
cor= 0.994
C145
UMI−based deduplication
Coordinate−based deduplication
cor= 0.964
C147
Coordinate−based deduplication
No deduplication
cor= 0.974
C147
UMI−based deduplication
No deduplication
cor= 0.988
C147
UMI−based deduplication
Coordinate−based deduplicaton
cor= 0.967
C148
Coordinate−based deduplication
No deduplication
cor= 0.966
C148
UMI−based deduplication
No deduplication
cor= 0.993
C148
UMI−based deduplication
Coordinate−based deduplication
cor= 0.954
W169
Coordinate−based deduplication
No deduplication
cor= 0.962
W169
UMI−based deduplication
No deduplication
cor= 0.995
W169
UMI−based deduplication
Coordinate−based deduplication
cor= 0.946

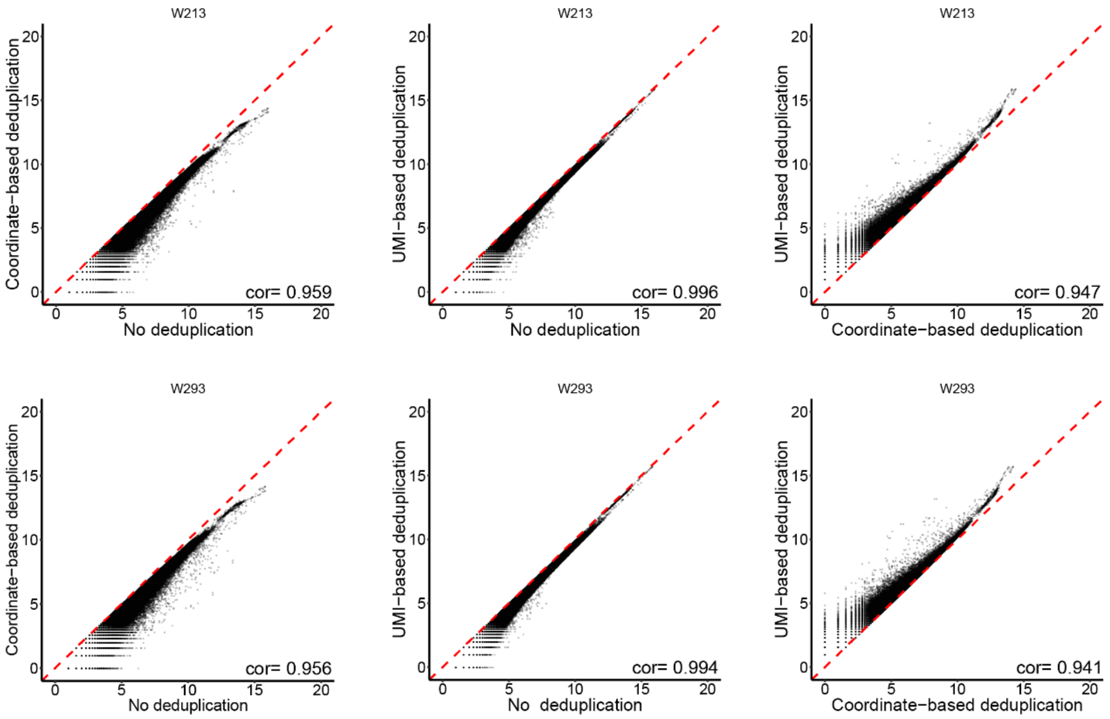


**Supplementary Figure 3: The correlations of quantification by removing PCR duplicates in different ways in different samples**

Each point is the number of Tn5 insertions in a 250 bp bin (log base 2). (a) Scatter plots contrasting quantification results by mapping coordinate-based deduplication (CD) and no deduplication (ND). (b) Scatter plots contrasting quantification result by UMI-based deduplication (UD) and ND. (c) Scatter plots contrasting quantification result by UD and CD.

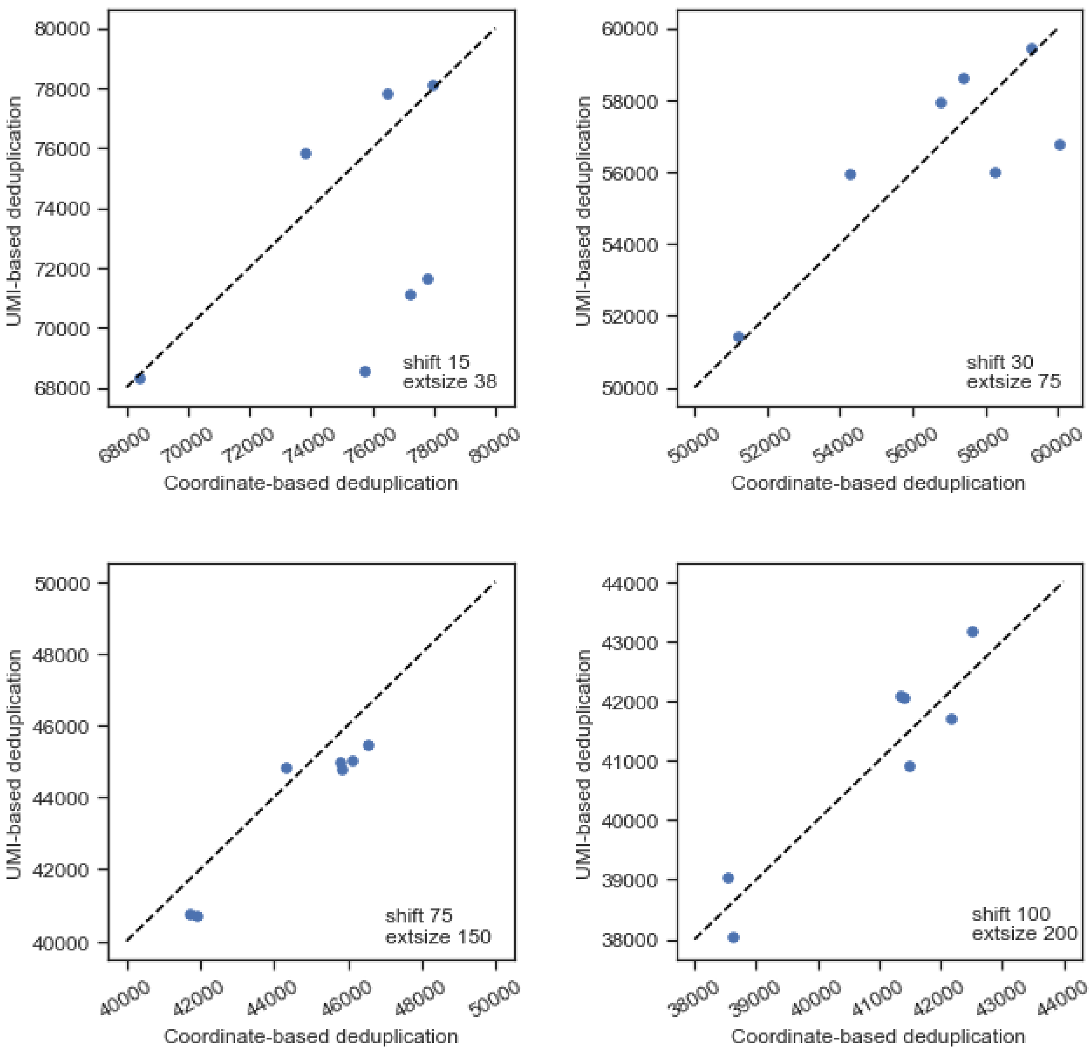


**Supplementary Figure 4: The comparisons of peak number generated by removing PCR duplicates in different ways in different samples.**

The parameters of calling peaks are showed in the lower right corner of each diagram.

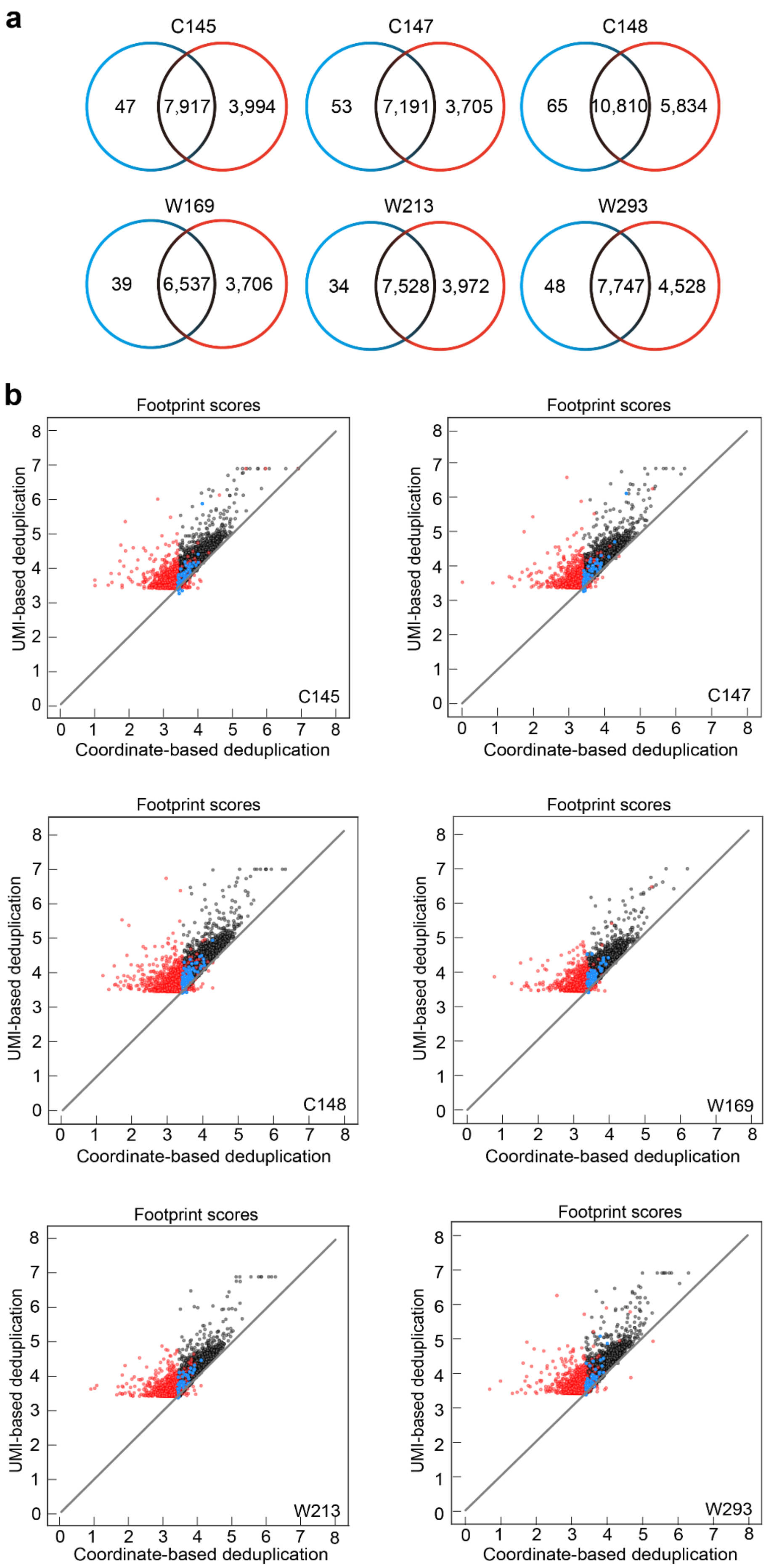
a
C145
47
7,917
3,994
C147
53
7,191
3,705
C148
65
10,810
5,834
W169
39
6,537
3,706
W213
34
7,528
3,972
W293
48
7,747
4,528
b
Footprint scores
UMI-based deduplication
Coordinate-based deduplication
C145
C147
C148
W169
W213
W293

**Supplementary Figure 5: The comparisons of footprints identified by removing PCR duplicates in different ways in different samples**

(a) Venn diagram displaying the footprints identified by pyDNase (Wellington Footprint *P-value* < $10^{-30}$) with CD (blue) and UD (red). The common footprints (black) are defined as those overlapped by at least one base. (b) The relationship between footprint scores calculated based on CD and UD in different samples. The footprint scores are calculated as $log(-log_{10}(P\text{-}value))$. The colors of points are the same as the Venn diagram in (a). The sample names are showed in the lower right corner of each diagram.

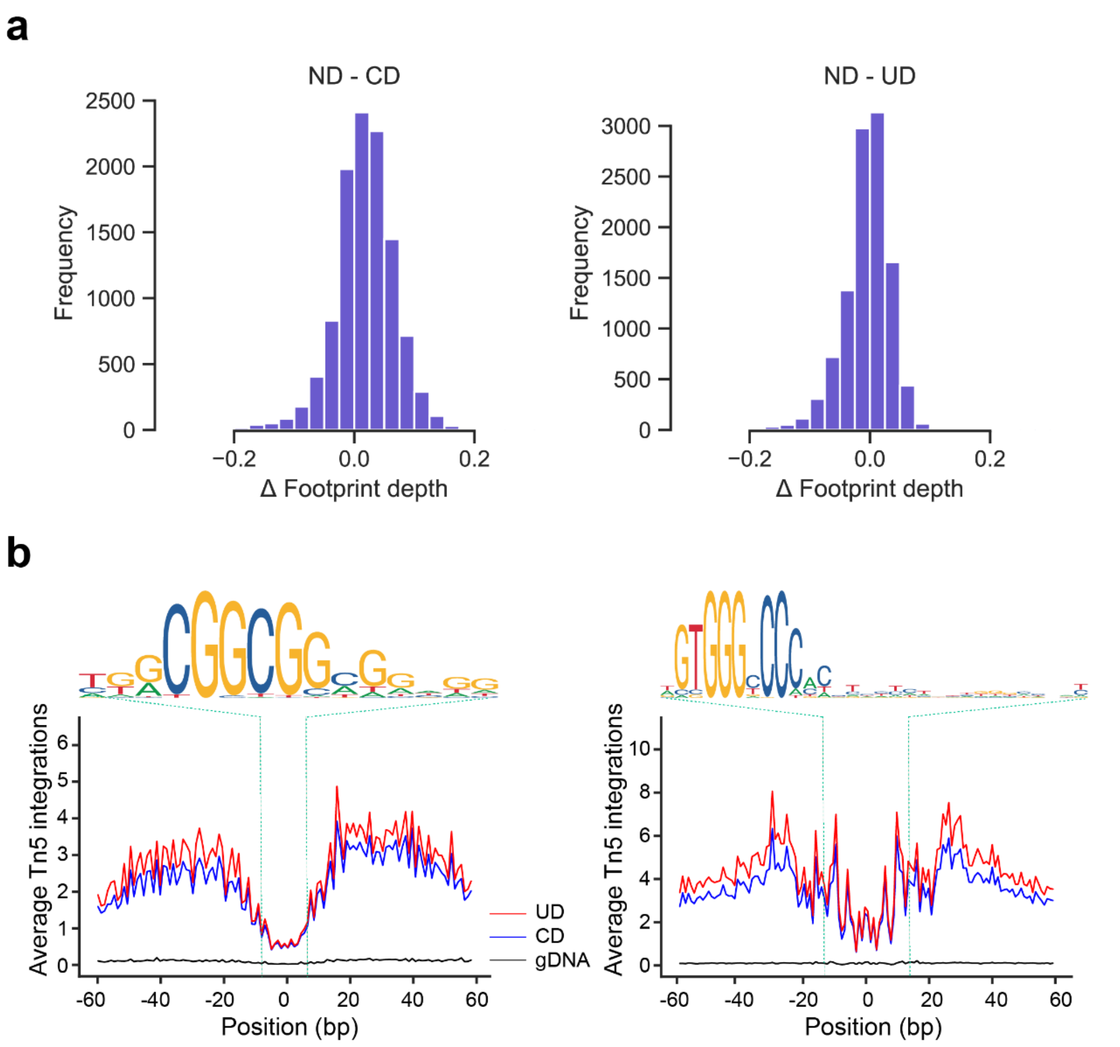


**Supplementary Figure 6: The comparisons of footprint pattern with removing PCR duplicates in different ways in sample C109.**

(a) The distribution of the differences in FPD values between ND and CD or ND and UD datasets. (b) Averaged Tn5 insertions per base for AP2/ERF (left) and TCP1(right) motif sites, which are overlapped with common pyDNase footprints identified in both CD and UD modes and located in the top 25% of the most accessible regions. The motifs are shown at the top of each diagram. Vertical lines (green) depict the edges of the motif match. The black line is genomic DNA sequencing data of Tn5-tagmented libraries (14.6-fold coverage).

**Supplementary Table 1: Primers used for UMI-ATAC-seq**

| Application | Primer name | Primer sequence |
|---|---|---|
| Tn5 assembly (ME sequence denotes in green; UMI sequence denotes in red; N denotes a random base) | MErev | [phos]CTGTCTCTTATACACATCT |
| | ME-A6 | CGACGCTCTTCCGATCTNNNNNNAGATGTGTATAAGAGACAG |
| | ME-A13 | CGACGCTCTTCCGATCTNNNNNNNNNNNNNAGATGTGTATAAGAGACAG |
| | ME-A20 | CGACGCTCTTCCGATCTNNNNNNNNNNNNNNNNNNNNAGATGTGTATAAGAGACAG |
| | ME-A25 | CGACGCTCTTCCGATCTNNNNNNNNNNNNNNNNNNNNCTGCTAGATGTGTATAAGAGACAG |
| | ME-B | GTCTCGTGGGCTCGGAGATGTGTATAAGAGACAG |
| PCR | PCR Primer 1 | AATGATACGGCGACCACCGAGATCTACACACACTCTTTCCCTACACGACGCTCTTCCGA |
| | Barcoded PCR Primer 2 (barcodes denote in red) | CAAGCAGAAGACGGCATACGAGATTGCCTCTTGTCTCGTGGGCTCGGAGATGT |
| | | CAAGCAGAAGACGGCATACGAGATTCCTCTACGTCTCGTGGGCTCGGAGATGT |
| | | CAAGCAGAAGACGGCATACGAGATATCACGACGTCTCGTGGGCTCGGAGATGT |
| | | CAAGCAGAAGACGGCATACGAGATACAGTGGTGTCTCGTGGGCTCGGAGATGT |
| | | CAAGCAGAAGACGGCATACGAGATCAGATCCAGTCTCGTGGGCTCGGAGATGT |
| | | CAAGCAGAAGACGGCATACGAGATACAAACGGGTCTCGTGGGCTCGGAGATGT |
| | | CAAGCAGAAGACGGCATACGAGATACCCAGCAGTCTCGTGGGCTCGGAGATGT |
| Sequence | Read1 Sequencing Primer/TrueSeq | ACACTCTTTCCCTACACGACGCTCTTCCGATCT |

**Supplementary Table 2: The summary statistics for UMI-ATAC-seq data**

| Sample | Total read pairs | Mapping rate | Coordinate-based duplication rate | UMI-based duplication rate | UMI rescue rate | Fraction of reads in peaks (FRiP) | TSS enrichment |
|---|---|---|---|---|---|---|---|
| C019 | 31,959,021 | 94.89% | 33.43% | 25.74% | 23.00% | 65.67% | 14.41 |
| C145 | 31,348,458 | 94.52% | 36.25% | 29.36% | 19.01% | 62.91% | 14.04 |
| C147 | 32,109,151 | 95.76% | 43.74% | 38.05% | 13.01% | 63.91% | 13.97 |
| C148 | 37,431,383 | 95.37% | 37.26% | 30.55% | 18.01% | 63.59% | 13.45 |
| W169 | 24,208,294 | 95.76% | 34.19% | 27.69% | 19.01% | 66.81% | 15.25 |
| W213 | 34,403,390 | 95.51% | 35.78% | 28.98% | 19.01% | 60.88% | 13.34 |
| W293 | 29,594,417 | 94.24% | 37.83% | 31.02% | 18.00% | 65.41% | 14.68 |
